# MEK1/2 inhibition decreases pro-inflammatory responses in macrophages from people with cystic fibrosis and mitigates severity of illness in experimental murine methicillin-resistant *Staphylococcus aureus* infection

**DOI:** 10.1101/2023.01.22.525092

**Authors:** Mithu De, Katherine B. Hisert, W. Conrad Liles, Anne M. Manicone, Emily A. Hemann, Matthew E. Long

**Affiliations:** Department of Internal Medicine, Division of Pulmonary, Critical Care, and Sleep Medicine, The Ohio State University, Columbus, OH; Department of Microbial Infection and Immunity, The Ohio State University, Columbus, OH; Department of Medicine, National Jewish Health, Denver, CO; Department of Medicine, Division of Infectious Diseases, University of Washington, Seattle, WA; Department of Medicine, Division of Pulmonary, Critical Care, and Sleep Medicine, University of Washington, Seattle, WA; Department of Medicine, Center for Lung Biology, University of Washington, Seattle, WA

**Author notes:** Corresponding Author: Matthew E. Long 473 W. 12^th^ Ave DHLRI Room 201, Columbus, OH 43210.

## Abstract

Chronic pulmonary bacterial infections and associated inflammation remain a cause of morbidity and mortality in people with cystic fibrosis (PwCF) despite new modulator therapies. Therapies targeting host factors that dampen detrimental inflammation without suppressing immune responses critical for controlling infections remain limited, while the acquisition of antibiotic resistance bacterial infections is an increasing global problem, and a significant challenge in CF. Pharmacological compounds targeting the mammalian MAPK proteins MEK1 and MEK2, referred to as MEK1/2 inhibitor compounds, have potential combined anti-microbial and anti-inflammatory effects. Here we examined the immunomodulatory properties of MEK1/2 inhibitor compounds PD0325901, trametinib, and CI-1040 on CF innate immune cells. Human CF macrophage and neutrophil phagocytic functions were assessed by quantifying phagocytosis of serum opsonized pHrodo red *E. coli*, *Staphylococcus aureus*, and zymosan bioparticles. MEK1/2 inhibitor compounds reduced CF macrophage pro-inflammatory cytokine production without impairing CF macrophage or neutrophil phagocytic abilities. Wild-type C57BL6/J and *Cftr*^tm1kth^ (F508del homozygous) mice were used to evaluate the in vivo therapeutic potential of PD0325901 compared to vehicle treatment in an intranasal methicillin-resistant *Staphylococcus aureus* (MRSA) infection with the community-acquired MRSA strain USA300. In both wild-type and CF mice, PD0325901 reduced infection related weight loss compared to vehicle treatment groups but did not impair clearance of bacteria in lung, liver, or spleen 1 day after infection. In summary, this study provides the first data evaluating the therapeutic potential of MEK1/2 inhibitor to modulate CF immune cells, and demonstrates that MEK1/2 inhibitors dampen pro-inflammatory responses without impairing host defense mechanisms mediating pathogen clearance.

## INTRODUCTION

Mutations in the cystic fibrosis transmembrane conductance regulator (CFTR) that impair this ion channel’s function result in the disease cystic fibrosis (CF). CF has many pathophysiological manifestations; however CF pulmonary disease remains the main driver of morbidity and mortality. Decades of investment in preclinical research have recently culminated in breakthrough new therapeutic highly effective modulator therapies (HEMT) that restore CFTR function and have rapidly changed the landscape of CF clinical disease and outcomes. However, despite the significant improvements in a majority of people with CF (PwCF) who have access to HEMT, established chronic bacterial colonization is not eradicated in many PwCF receiving modulator therapy, and is anticipated to remain a healthcare challenge for PwCF^1^. In addition, there remains a significant minority of PwCF who are not eligible for current modulator therapies in whom improved therapies to mitigate lung damage caused by chronic airway infection and inflammation are still needed. Novel strategies to alleviate pulmonary inflammation and infection are essential to address these gaps in care.

Innate immune cells, including monocytes, macrophage, and neutrophils, that reside in the lung or are recruited during infection or injury are critical for eliminating infections and resolving the inflammatory response. However, the accumulation of inflammatory cells in the CF lung, especially neutrophils, contributes to tissue pathology^2^, and therapeutic approaches to reduce the tissue damaging effects of innate immune cells in the CF lung have been elusive. Our previous studies and others’ have demonstrated that targeted inhibition of mammalian protein kinases MEK1/2 is a promising therapeutic strategy to reduce detrimental inflammation during the context of lung injury and infection^3, 4, 5^. Development of small molecules to inhibit the MEK1/2-ERK1/2 pathway has taken place over the last several decades due to the high levels of over-activation of this pathway in several types of human cancer^6^. Four different MEK1/2 inhibitor compounds are currently FDA approved, with several additional compounds under investigation in clinical trials^7^. Furthermore, novel therapeutic applications of MEK1/2 inhibitor compounds, including administration during respiratory viral infections, is an active area of preclinical and translational research^8, 9, 10^. The current study was undertaken to determine whether MEK1/2 inhibitor compounds can reduce phagocyte-mediated inflammation in CF without impairing phagocyte host defense mechanisms. This study utilized primary human blood-derived macrophages and neutrophils from PwCF to examine how administration of MEK1/2 inhibitor compounds modulate CF macrophage pro-inflammatory responses to LPS and CF macrophage and neutrophil phagocytic abilities. Wild-type C57BL6/J and *Cftr*^tm1kth^ (F508del homozygous) mice were used to evaluate if delivery of a MEK1/2 inhibitor impaired host defense following infection with methicillin-resistant *Staphylococcus aureus* (MRSA).

## METHODS

### Reagents

The MEK1/2 inhibitor compounds PD0325901 (catalog # S1036), CI-1040 (catalog # S1020), and trametinib (catalog # S2673) were purchased from Selleck for use in in vitro cell culture experiments. PD0325901 and trametinib were used at 0.5 μM and CI-1040 was used at 10 μM. PD0325901 (catalog # PZ0162) used for in vivo experiments was from Sigma. Recombinant human M-CSF was from Peprotech (catalog # 300-25). Ficoll Paque Plus (catalog # 17-1440-03) was from Cytiva. Human AB serum was from Sigma (catalog # H6914). LPS from *Pseudomonas aeruginosa* was from Sigma (catalog # L8643).The pHrodo Red *E. coli* (catalog # P35361), *S. aureus* (catalog # A10010), and zymosan (catalog # P35364) bioparticles were from ThermoFisher Scientific. Antibodies used in these studies are listed in Supplemental Table 1.

### Animal Ethics and Mouse Infection

The Institutional Animal Care and Use Committee at The Ohio State University approved all studies (OSU IACUC #2020A00000081). *Cftr*^tm1kth^ F508del homozygous mice were acquired from the Case Western University Cystic Fibrosis Mouse Model Core. Mice were maintained ad libitum on golyteyl drinking water (25 mM sodium chloride, 10 mM potassium chloride, 40 mM sodium sulfate, 20 mM sodium bicarbonate, 18 mM PEG 3350). C57BL/6J wild-type mice were obtained from Jackson Laboratories. For infection, an overnight culture of methicillin-resistant *S. aureus* USA300 grown in BHI was sub-cultured to mid-logarithmic phase. Bacteria were washed twice with sterile PBS and diluted so that 1 × 10^7^ colony forming units (CFU) were delivered in a 50 μl inoculum. Mice were anesthetized by i.p. injection of ketamine/xylazine. Anesthetized mice were provided i.p. injection of sterile PBS containing the MEK1/2 inhibitor PD0325901 (20 mg/kg) or vehicle (DMSO) immediately prior to infection by intranasal delivery of the 50 μl inoculum. Animals were monitored for recovery from anesthesia and weight loss was measured daily. CF mice were sacrificed 24 hours after infection and lung, liver, and spleens were collected, homogenized in 1 ml of sterile PBS with the Precellys lysing kit beads (catalog # P000912-LYSK1-A), and serial dilutions of tissue homogenates were plated on TSA and incubated overnight at 37°C to allow colony growth. Colonies were enumerated and the total CFU recovered were calculated and normalized per gram of tissue collected.

### Cell culture

Human subjects research was approved by the Institutional Review Board at Nationwide Children’s Hospital and The Ohio State University. Blood donors were consented at Nationwide Children’s Hospital by clinical research coordinators supported by the Cure CF Columbus Research Development Program. Human blood was collected into EDTA-vacutainer tubes following approved IRB protocols and samples were de-identified. Peripheral blood mononuclear cells (PMBC) were isolated by Ficoll gradient centrifugation and differentiated to monocyte-derived macrophages for 7-10 days by incubation with 20 ng/ml recombinant human M-CSF. Macrophages were seeded into 24-well tissue culture plates at a density of 300,000 cells/well in HEPES-buffered RPMI-1640 containing 10% HI-FBS, penicillin/streptomycin, L-glutamine. Neutrophils were also isolated from peripheral blood by Ficoll gradient centrifugation and used immediately following isolation^11^.

### Phagocytosis of pHrodo Bioparticles

For opsonization of pHrodo labeled bioparticles, a frozen vial was resuspended in sterile HBSS containing calcium and magnesium (according to manufacturer’s recommendations) and placed in a water bath sonicator for 15 minutes to fully disperse particles. To opsonize the bioparticles, an equal volume of human serum was added to the particles and were incubated for 30 minutes at 37°C. Following opsonization, bioparticles were added to macrophage or neutrophil cultures at a concentration of 0.08 mg/ml. Neutrophil cultures were gently tumbled for 3 minutes at 37°C to ensure proper mixing followed by subsequent static incubation. Following a 1 hour incubation at 37°C, supernatants were aspirated and cells were washed twice with PBS to remove non-ingested particles. Macrophages were collected by addition of PBS containing 2 mM EDTA to the cultures and cells were gently lifted using a cell scraper, washed with cell staining buffer (catalog # 420201, BioLegend), and stored on ice and protected from light^12^. Data were collected on a Becton Dickinson LSRFortessa flow cytometer and analyses performed with FlowJo software. Cytochalasin D was added to samples at 2 μM as a control for inhibition of phagocytosis and was used to draw flow gates to identify the percent of cells negative and positive for pHrodo red.

### Macrophage LPS Stimulation for Western Blot and ELISA

Protein lysates were prepared as previously described^3^. In brief, RIPA lysis buffer containing protease and phosphatase inhibitors (Life Technologies) were added to cells on ice for 20 minutes. Cell lysates were scraped from the wells, collected into pre-chilled centrifuge tubes and the lysates were cleared by centrifugation at 14,000 × *g* for 5 minutes at 4°C. Quantitation of protein lysates was performed using a BCA assay (Life Technologies), and samples were prepared in order to load 5-10 ug of total protein per lane of a gel. Supernatants from stimulated cells were collected and centrifuged at 14,000 × *g* for 5 minutes to clear cellular debris and were stored at −80°C prior to ELISA analysis. Human IL-8/CXCL8 and TNF DuoSet ELISA (R&D Systems) were used to quantify IL-8 and TNF in supernatants.

## RESULTS

### Inhibition of the MEK1/2 Pathway Decreases CF Macrophage LPS-Induced IL-1β and IL-8 Production

Production and secretion of IL-1β and IL-8 are thought to be key drivers of inflammation in CF^13^. To test the hypothesis that MEK1/2 inhibitor compounds would reduce production of pro-IL1β and secretion of IL-8, CF macrophages were stimulated with *P. aeruginosa* LPS in media containing vehicle or one of the MEK1/2 inhibitor compounds, PD0325901, Trametinib, or CI-1040. When analyzed 4 hours after stimulation, addition of the MEK1/2 inhibitor compounds reduced MEK1/2-ERK1/2 pathway activation, as measured by reduced ERK1/2(pT202/pT204) (Fig. 1A), and the LPS-induced production of pro-IL-1β was also significantly reduced by addition of a MEK1/2 inhibitor compound compared to vehicle-treated control (Fig. 1A-B). Further, the LPS-induced secretion of IL-8 (Fig. 1C) and TNF (Fig. 1D) were significantly reduced by addition of the MEK1/2 inhibitor compounds compared to vehicle-treated samples. Together, these results demonstrate that MEK1/2 inhibitors are able to significantly reduce CF macrophage LPS-induction of pro-inflammatory cytokines associated with neutrophil recruitment and tissue damage.

**Figure 1.**
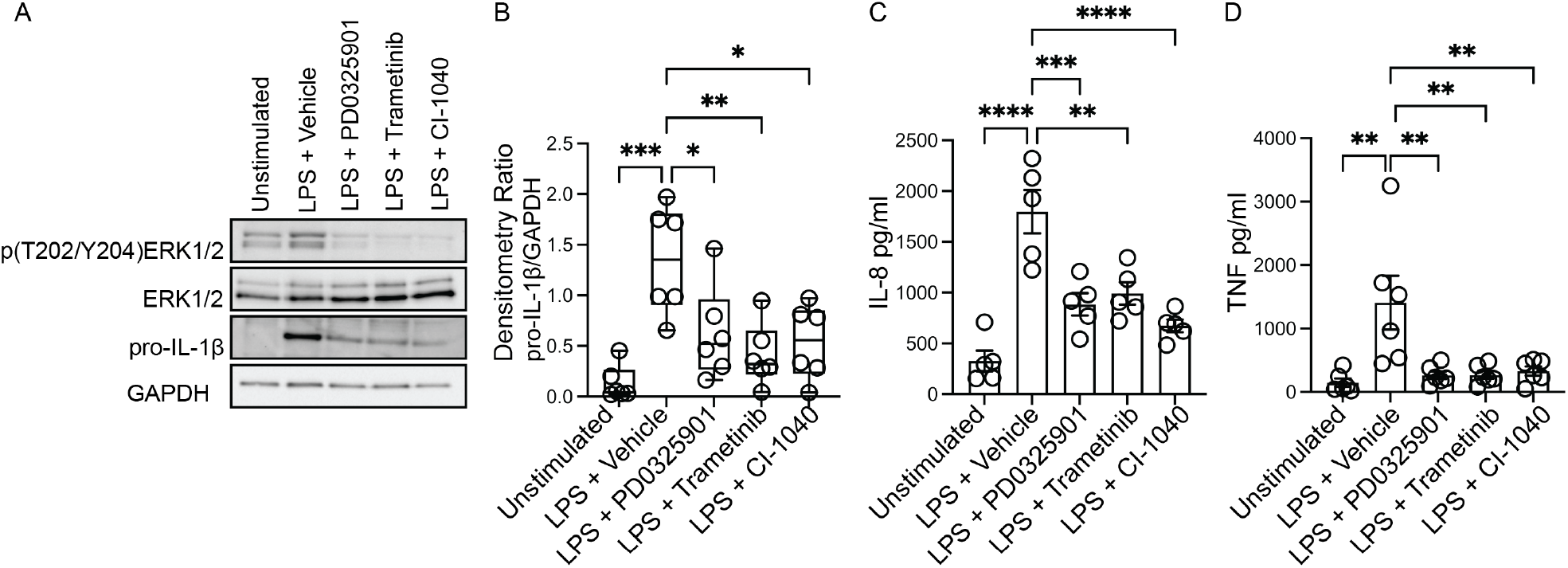
MEK1/2 inhibitors reduce CF macrophage LPS-induced pro-inflammatory responses. Human CF macrophage were stimulated with 50 ng/ml of *P. aeruginosa* LPS for 4 hours. (A) Protein lysates from a representative experiment from one donor, (B) densitometry quantitation of the ratio of pro-IL-1B to GAPDH from n=6 donors.(C) Measurement of the levels of IL-8 or (D) TNF from supernatants collected at 4 hours from n=5-6 donors. (B-D) Data are the mean ± SEM, each point represents one individual donor. Statistical analyses were performed by One-Way ANOVA with Tukey multiple comparisons, * *P*<0.05, ** *P*<0.01, *** *P*<0.001, **** *P*<0.0001.

### MEK1/2 Inhibitors Do Not Impair CF Macrophage or Neutrophil Phagocytosis

While production and secretion of cytokine and chemokines by macrophages modulate the inflammatory response, phagocytosis and phagosome acidification are critical cellular functions of both macrophage and neutrophils involved in host defense and homeostasis. To determine if MEK1/2 inhibitors reduced the ability of macrophage to perform these functions on gram-negative, gram-positive, or fungal microbes, phagocytosis and phagosome acidification were assayed with serum-opsonized pHrodo-labeled *E. coli* (Fig. 2A,D), *S. aureus* (Fig. 2B,E), or zymosan (Fig. 2C,F) bioparticles. The pHrodo label requires acidification of the phagosome compartment to increase fluorescence, therefore detection in this assay is dependent on both bioparticle internalization and phagosome maturation. The addition of cytochalasin-D during the phagocytosis assay was used as a control and significantly reduced and prevented bioparticle ingestion, in contrast to vehicle-treated macrophage, which exhibited robust phagocytic abilities as measured by the percent of macrophages positive for pHrodo-red labeling (Fig. 2 A-F). Macrophages treated with PD0325901, Trametinib, or CI-1040 did not have significantly reduced phagocytosis of either opsonized *E. coli* or opsonized *S. aureus* bioparticles compared to vehicle-treated controls. In addition, the ability of CF neutrophils to phagocytose serum-opsonized *E. coli* (Fig. 3A) or *S. aureus* (Fig. 3B) pHrodo bioparticles was not impacted by addition of MEK1/2 inhibitors. Together, these results indicate inhibition of MEK1/2 does not impact human macrophage or neutrophil phagocytosis or phagosome acidification of opsonized microbes.

**Figure 2.**
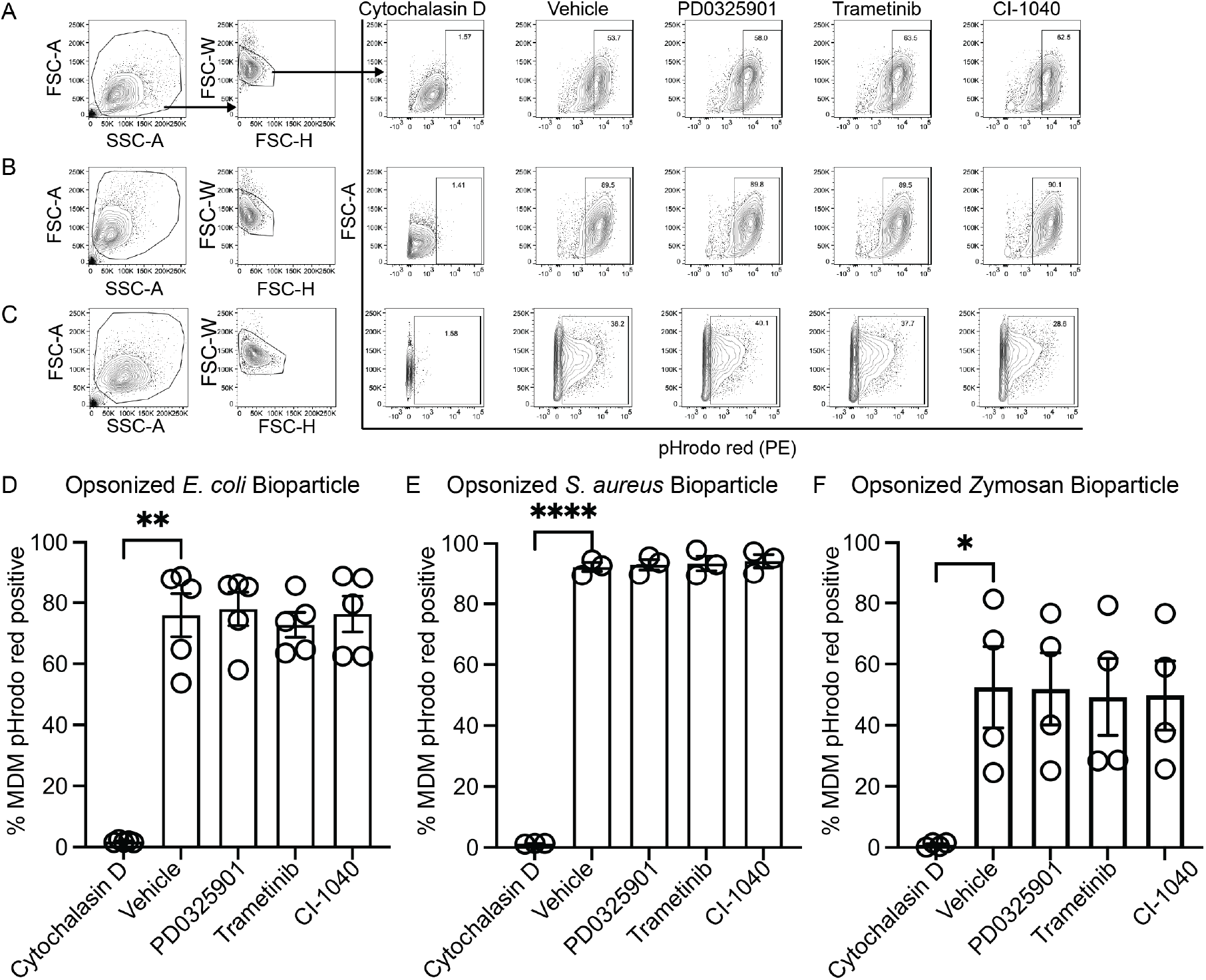
MEK1/2 inhibitors do not impair CF macrophage phagocytosis. Representative flow cytometry gating used for phagocytosis of (A) pHrodo Red *E. coli* bioparticles, (B) pHrodo Red *S. aureus* bioparticles, or (C) pHrodo Red Zymosan bioparticles. Quantitation of the percent of macrophages positive for pHrodo red fluorescence following incubation with serum opsonized (D) pHrodo Red *E. coli* bioparticles, (E) pHrodo Red *S. aureus* bioparticles, or (F) pHrodo Red Zymosan bioparticles. Data are the mean ± SEM and each point represents data from one individual donor. Statistical analyses were performed by One-Way ANOVA with Tukey multiple comparisons, * *P*<0.05, ** *P*<0.01, **** *P*<0.0001.

**Figure 3.**
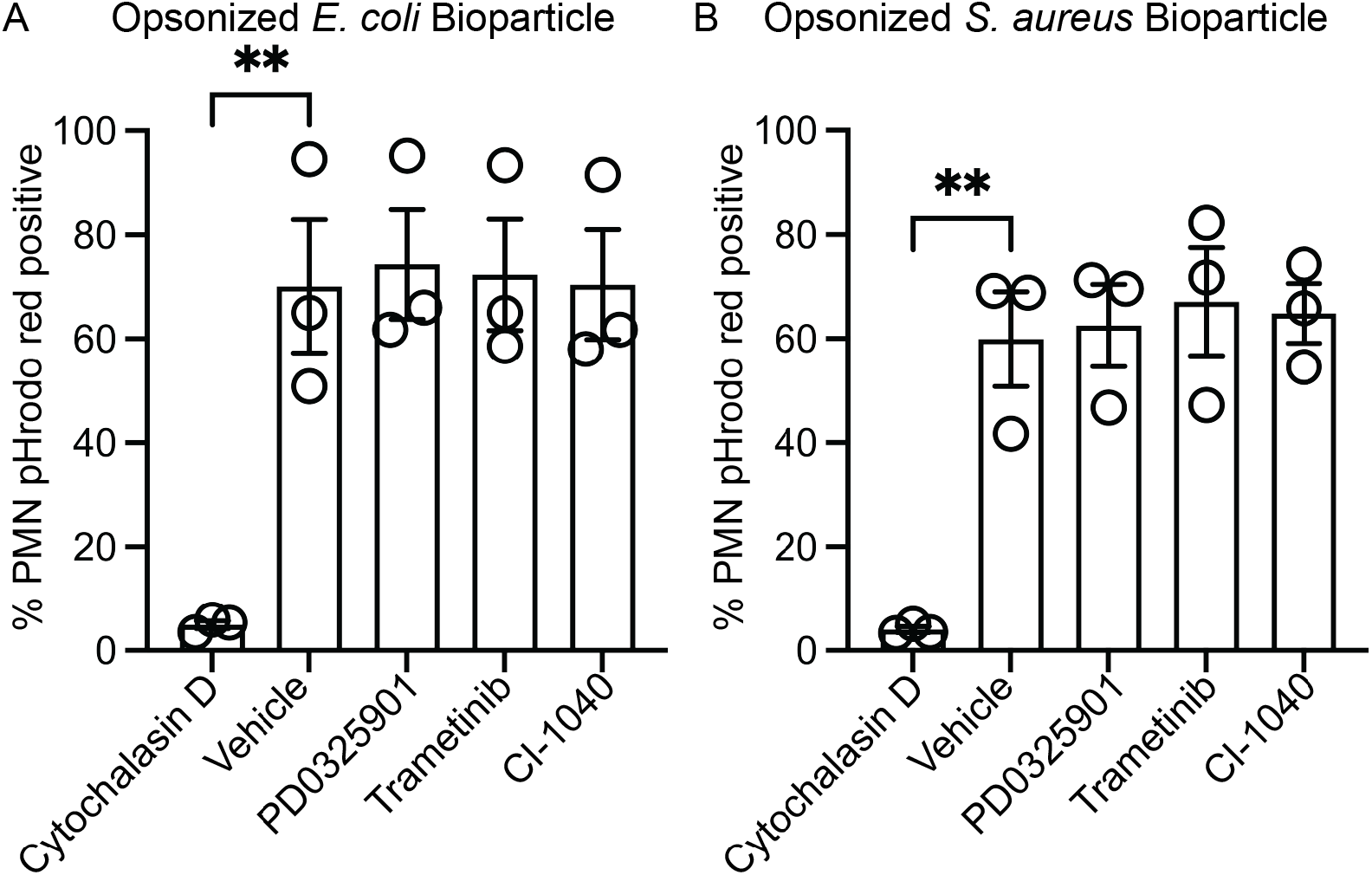
MEK1/2 inhibitors do not impair CF neutrophil phagocytosis. Quantitation of the percent of neutrophils positive for pHrodo red fluorescence following incubation with opsonized (A) pHrodo red *E. coli* bioparticles or (B) pHrodo red *S. aureus* bioparticles. Data are the mean ± SEM and each point represents one individual donor. Statistical analyses were performed by One-Way ANOVA with Tukey multiple comparisons** *P*<0.01.

### *In vivo* MEK1/2 Inhibitor Delivery Does Not Impair Host Defense During MRSA Infection in CF Mice

Inhibition of innate immune inflammatory functions may decrease over- exuberant inflammation, but could result in impaired host defense response during *in vivo* infections^14, 15^. We thus sought to interrogate whether the anti-inflammatory effects of a MEK1/2 inhibitor compound would increase bacterial replication or dissemination *in vivo*. Mice were treated i.p. with either vehicle or PD0325901 immediately prior to intranasal infection with the MRSA strain USA300. At 24 hours after infection, wild-type mice receiving PD0325901-treatment had significantly less weight loss, an indicator of illness, compared to vehicle-treated animals, which was sustained through day 4 of infection (Fig. 4A); all mice had returned to similar levels of weight as mock control on days 5-6 after infection. We next utilized the CF mouse model to determine if PD0325901 treatment also alleviated weight loss as a marker of illness without impairing bacterial clearance. Similar to findings with wild-type mice, CF mice receiving PD0325901 treatment experienced significantly less weight loss compared to vehicle treated animals at day 1 after infection (Fig. 4B). At this 1 day timepoint in CF mice, CFU were measured in lung, liver, and spleen homogenates to quantify the bacterial burdens; there were no significant differences in bacterial clearance in the PD0325901 treated group compared to vehicle treated controls (Fig. 4C). Together, these results demonstrate that MEK1/2 inhibitor-treatment reduced a key marker of illness following infection but did not impair bacterial clearance.

**Figure 4.**
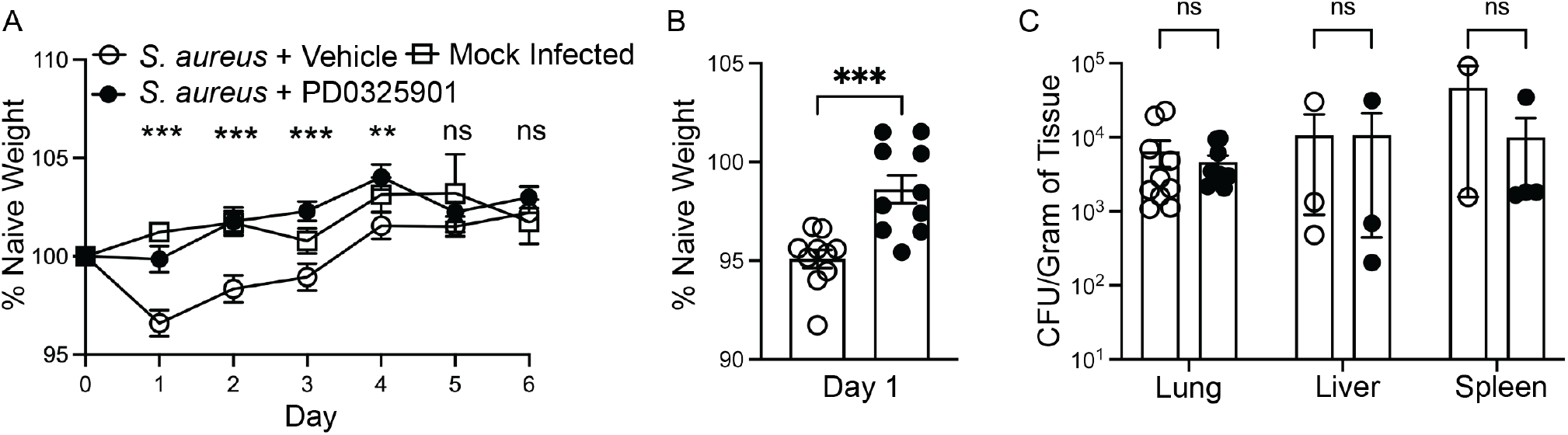
MEK1/2 inhibitor administration does not impair host defense of CF mice during *S. aureus* infection. Anesthetized mice were provided i.p. treatment with 20 mg/kg PD0325901 (black filled circles) or vehicle control (open circles) immediately prior to intranasal inoculation with 1 × 10^7^ CFU of *S. aureus* USA300. (A) Weight loss of C57BL6/J mice was measured following infection or mock infection for six days. Data are the mean ± SEM from n=5 males and n=5 females for each infected treatment group, and n=4 total for mock infected. (B) Weight loss of *Cftr*^tm1kth^ F508del homozygous mice following infection. Data points represent an individual mouse and are combined from three independent experiments, n=5 males and n=5 females for each treatment group. (C) Tissue homogenates from lung, liver, and spleen were used to enumerate total colony forming units (CFU) and normalized per gram of tissue collected. Each data point is from an individual mouse; data points and were omitted from the graphs if the CFU recovery was below the limit of detection (99 CFU/ml in a single spot). Statistical analysis were performed with (A) Two-Way ANOVA with multiple comparisons, (B) Unpaired t-test, and (C) multiple unpaired t-tests; \*\**P*<0.01, \*\*\**P*<0.001, ns not significant.

## DISCUSSION

Despite the fact that HEMT has significantly improved clinical outcomes and health in many PwCF, there is still a critical need for novel therapies for people with CF with established lung disease associated with chronic bacterial infection and inflammation. Access to HEMT therapy remains restricted based on eligibility of only specific *CFTR* mutations, insurance and socioeconomic status impact access to HEMT, and HEMT is still not available to PwCF in many countries. In addition, chronic infections remain challenges in CF, even for PwCF on HEMT. According to patient registry data, in the USA the prevalence of *S. aureus* infections in PwCF has been over 60% since 2006^16^. Due to early colonization and high prevalence of respiratory infections, acquired and intrinsic antimicrobial resistance of airway bacteria are straining our arsenal of therapeutic options. Novel therapeutic strategies to reduce detrimental inflammatory responses while combatting infection have major potential to slow pathology in CF, reducing symptom burden and enhancing patient longevity.

The unexpected clinical trial results for a previously developed anti-inflammatory therapy for PwCF highlighted that suppressing specific arms of the inflammatory response can lead to impaired host defenses, increased growth of *P. aeruginosa*, and increased pulmonary exacerbations^14, 15^. To address these concerns, experimental anti-inflammatory therapies need to undergo extensive and rigorous preclinical studies to investigate the potential for impaired host defense. Our previous studies using non-CF human macrophages and wild-type mice demonstrated that MEK1/2 inhibitors modulated macrophage polarization *in vitro* and *in vivo*, and mitigated illness in wild-type mice when therapeutically delivered after initiation of experimental LPS-induced acute lung injury or *P. aeruginosa* pneumonia^3, 4^. Other non-CF models of infection and inflammation demonstrated complementary findings to our results, supporting the anti-inflammatory therapeutic potential of MEK1/2 inhibitors^5, 17, 18, 19, 20^. However, to date there have been no studies investigating the effects of MEK1/2 inhibitors on CF immune cells, and the application of a MEK1/2 inhibitor in preclinical models of pulmonary MRSA infection has not been reported. This study is the first to investigate the therapeutic potential and immunomodulatory roles of MEK1/2 inhibitors using human CF macrophages and neutrophils, and in an experimental murine MRSA infection model in wild-type and CF mice. The results from this study demonstrated that inhibition of the MEK1/2 pathway decreased LPS-induced production of IL-1β and secretion of IL-8 by CF macrophages, two key inflammatory mediators that potentiate tissue damage and recruitment of neutrophils. In addition, the data presented here indicate that inhibition of the MEK1/2 pathway does not decrease the phagocytic abilities of CF macrophages and neutrophils or affect phagosome acidification. Finally, the data demonstrated that administration of a MEK1/2 inhibitor compound to mice at the time of MRSA infection did not impair bacterial clearance or result in increased dissemination of infection. Instead, groups of mice treated with the MEK1/2 inhibitor had reduced weight loss as a marker of illness compared to vehicle-treated groups. These new findings are consistent with previous reports demonstrating that MEK1/2-ERK1/2 pathway inhibition reduces airway epithelial cell, including CF epithelial cell, production of IL-8^21, 22^, and that inhibition of the MEK1/2-ERK1/2 pathway can prevent airway epithelial cell CFTR degradation^23, 24^. Combined, these new data support the hypothesis that MEK1/2 inhibitor compounds exert anti-inflammatory effects without impairing host defense mechanisms, and thus may have a high potential as therapies for PwCF.

There are several limitations of the current study. First, while the CF mouse model utilized recapitulates many phenotypes of CF, the mouse does not reproduce all of the aspects of human CF pulmonary disease, such as spontaneous airway disease or have airway acidification defects. In addition, the MRSA infection employed results in a modest and transient disease, which does not fully capture the chronic infections commonly found in CF. However, this murine model is still an important tool for the development of preclinical studies as it models complex immune cell interactions with bacteria in the lung. Second, while our approach utilized the MRSA USA300 strain, which is associated with community-acquired human disease, CF clinical isolates of *S. aureus* may have significant genetic and phenotypic differences to produce different inflammatory host responses. For example, infection with *S. aureus* small colony variants (SCV) have been demonstrated to elicit worse outcomes in PwCF and infection induces an increased inflammatory response compared to non-SCV *S. aureus* infection in the CF rat^25, 26^. Overall, the current findings support a rationale for future investigative preclinical studies utilizing CF models, such as the CF rat^25, 27^, that more faithfully recapitulate CF pulmonary disease in conjunction with chronic infection or poly-microbial infection.

The translational potential of this work is highly relevant, as a recent human clinical trial (NCT04776044) has tested the therapeutic potential of the MEK1/2 inhibitor compound ATR-002 in the context of SARS-CoV-2 infection^10^. Future human clinical trials with ATR-002 may provide additional data to help determine the safety and efficacy of this compound during human respiratory infections, and could serve as a foundation for evaluating the safety for translational application in PwCF. Significantly for CF, there is evidence that the ATR-002 compound has direct antibacterial effects on gram-positive organisms, including *S. aureus^28^*. Future preclinical studies for CF should be designed to harness the combined antimicrobial and potential anti-inflammatory effects of this MEK1/2 inhibitor compound. In summary, this study provides the first CF data evaluating the immunomodulatory and therapeutic potential of MEK1/2 inhibitor compounds. The findings support the hypothesis that inhibition of the MEK1/2 pathway is a therapeutic target to reduce inflammation without impairing cellular and organism level host defense mechanisms. Future preclinical studies to provide additional rigorous assessment of the impact on inflammation, host defense, and the antimicrobial potential of MEK1/2 inhibitor compounds should be performed with the goal to developing a path to human CF translational studies.

## ACKNOWLEDGEMENTS

This work was supported in part by the Cure CF Columbus Translational Core (C3TC). C3TC is supported by the Division of Pediatric Pulmonary Medicine, the Biopathology Center Core, and the Data Collaboration Team at Nationwide Children’s Hospital. Grant support was provided by The Ohio State University Center for Clinical and Translational Science (National Center for Advancing Translational Sciences Grant UL1TR002733, the Cystic Fibrosis Foundation Research Development Programs MCCOY19RO and SINGH15RO, Cystic Fibrosis Foundation awards LONG19F5-CI and LONG21R3 to M.E.L., and MANICO19G0 to A.M.M, and the National Institutes of Health awards K08 HL136786 to K.B.H, K22 AI146141 to E.A.H. and R01 HL144656 to A.M.M. This research was supported by the Flow Cytometry Shared Resource at The Ohio State University supported by NCI P30CA16058.

---

**Supplemental Table 1:** List of antibodies used for western blot.

| Target | Company | Species | Catalog # |
| --- | --- | --- | --- |
| GAPDH (14C10) | Cell Signaling | Rabbit | 2118 |
| P44/42 MAPK (ERK1/2) (137F5) | Cell Signaling | Rabbit | 4695 |
| p-P44/42 MAPK (ERK1/2) (T202/Y204) | Cell Signaling | Rabbit | 9101 |
| IL-1β (D3U3E) | Cell Signaling | Rabbit | 12703 |

